# Surviving in the mountains: Temperature and elevation have contrasting physiological effects on the hoverfly *Eristalis tenax* in the Himalayas

**DOI:** 10.1101/2024.12.14.628473

**Authors:** Gauri Gharpure, Jagath Vedamurthy, Sakshi Priya, Geetha G Thimmegowda, Shannon B. Olsson

## Abstract

Insect populations are experiencing a global decline due to a variety of human-linked environmental changes. Among these changes, how insects’ physiology might be affected by predicted upslope migration due to climate change is unknown. Being ectotherms, insect physiology is impacted by abiotic factors like ambient temperature that change with elevation. Here, we performed *in situ* experiments to assess the sensory and cardiac physiology of an important generalist pollinating hoverfly *Eristalis tenax* (Diptera: Syrphidae), across different elevations in the eco-sensitive and biodiverse Himalayan mountains. We built a portable physiology setup and measured hoverfly antennal responses towards common floral volatiles at 3600 masl and 4200 masl. We also recorded their heart rate at 3000 masl, 3500 masl and 4000 masl. We report the first *in situ* physiology experiments performed in the high-altitude Himalayas. Our results show a contrasting impact of elevation and temperature on the sensory and cardiac physiology of hoverflies, with antennal sensitivity decreasing with increasing elevation, while average heart rate increased with temperature, independent of elevation. With upslope migration and climate warming, consequent sensory mismatches and cardiac stress could have deleterious effects on the health of both hoverflies and the vulnerable Himalayan ecosystem.

## INTRODUCTION

Montane ecosystems span nearly 12% of the Earth’s surface (Tito et al., 2020) and are witness to the effects of rapid environmental changes due to anthropogenic impacts such as land development, environmental degradation, biodiversity loss, agricultural displacement, and invasive species. Resident montane organisms have been known to be severely affected, leading to declines in species abundance, composition and upslope migration (IPCC 2019 Technical Summary). Studies on the impacts of climate warming on species have overwhelmingly been performed in temperate mountains, leaving the fate of tropical mountains, like the Himalayas, unclear (Chen and Lewis, 2023). The Himalayan mountains constitute an important ecologically sensitive region with high biodiversity and endemism (Brooks et al., 2006). Yet, research about the biodiversity from this region does not reflect this species richness, especially for insects, the biggest multicellular taxon (Rana et al., 2021).

Insects perform several functions in the ecosystem, with pollination being one of the most well-known ecosystem services (Peña-Kairath et al., 2023; Stephens et al., 2023). Wild insect pollinators constitute important nodes in ecological networks, often surpassing domesticated bees in terms of pollination capacity (Garibaldi et al., 2013). Previous studies have shown that insects, along with other taxa, have been shifting their range upwards due to climate warming at lower elevations (McCain and Garfinkel, 2021). These displaced organisms are often restricted by the low atmospheric oxygen at higher elevations due to low atmospheric pressure (Dillon et al., 2006; Spence and Tingley, 2020). How these ecologically important insects are affected by the functional hypoxia, low humidity, and altered nutrient cycling at higher elevations remains understudied in montane ecosystems (Harrison et al., 2018; Shah et al., 2020), particularly in the Eastern Himalayas (Trunschke et al., 2024).

Previous studies concerning the survival of insects in different environmental conditions have focused on either community-level traits like network analysis, phenology studies, (Hassall et al., 2019; Gérard et al., 2020; Spence and Tingley, 2020; Johnson et al., 2023), or species-level traits like critical thermal maxima (Sunday et al., 2011; Khelifa et al., 2019). A small number of studies have also assessed insect pollinators in the Himalayas, such as thermal melanism (Gautam and Kunte, 2020), pollinator assemblages (Gurung et al., 2018, Basnett et al., 2024) and plant-pollinator networks (Rather et al., 2023). Physiological experiments on insects have primarily been performed with lab-reared populations, or with individuals brought back to lab. However, this fails to capture the physiological status of insects directly exposed to environmental changes.

Being poikilotherms, insects’ metabolism is significantly affected by external abiotic factors, making them especially vulnerable to changes in environmental conditions. Average heart rate can be a good proxy for their metabolic rate. Insect heart rate has been known to be affected by extrinsic factors like pollution levels (Thimmegowda et al., 2020), diet (Bazzell et al., 2013), nutritional status (Ellison et al., 2015; Gill et al., 2015) and temperature (Richards, 1963; Lagerspetz and Perttunen, 1962; Andersen et al., 2015), but less is known about how cardiac physiology might be affected by functional hypoxia across elevations. Further, the sensory perception and behaviour of insects towards floral resources (Gérard et al., 2023), as well as neurophysiological parameters (Andersen et al., 2018; Andersen et al., 2023) can also be impacted by climate and environmental change. There have been a few attempts to understand electrophysiological responses of insects in field through portable electroantennograms using excised antennae of different moth species (Pawson et al., 2020; Terutsuki et al., 2021; Schott et al., 2013; Mateus 2016). However, *in situ* whole-body preparations that reflect current physiological states of insects in their local environment have not been performed. Together, these issues have resulted in a dearth of field-based studies to understand the fate of insect-mediated plant-pollinator interactions in light of climate change (Hegland et al., 2009; Trunschke et al., 2024).

Dipterans are the major group of pollinators in mid-elevations of the Himalayas (Gurung et al., 2018; Basnett et al., 2024), and especially hoverflies, globally known to be important pollinators (Rader et al., 2020; Raguso, 2020). One such hoverfly species is *Eristalis tenax* (Syrphidae: Diptera), a generalist pollinator found across the globe (Klecka et al., 2018; Lucas et al., 2018; Doyle et al., 2023). It occurs at mid-high elevations in the Himalayas and pollinates both wildflowers and crops in the region (Sengupta et al., 2016; Nordström et al., 2017; Sinha et al., 2022; Rather et al., 2023). It is known to be migratory in Australia, Europe, North America, and Asia (Doyle et al., 2023), therefore contributing to both short and long-distance pollination (Wotton et al., 2019). Here, we performed *in situ* experiments on wild-caught individuals of *Eristalis tenax* at different elevations from 3000 – 4200 masl in the Western (Spiti valley, Himachal Pradesh) and Eastern (Lachen valley, Sikkim) Himalayas to assess the impact of environmental change on the sensory and cardio-physiology of an important insect pollinator. The resulting data not only helps us to understand the current survival of these Syrphid flies, but also predict how their populations might respond to environmental changes in the future (Shah et al., 2020).

## METHODS

### Study sites

The measurement of antennal sensitivity was performed at two elevations in western Himalayas (Spiti Valley, Himachal Pradesh: Kaza (3600masl), Kibber (4200masl)) in June 2023. Heart rate recordings were performed at three elevations in eastern Himalayas (Lachen valley, Sikkim: Zima-III (3000masl), Thangu (3500masl), Chopta (4000masl)) in May and September 2022, and also in the same sites in Spiti valley as above. Control laboratory experiments were performed in Bengaluru (900masl) during January-March 2023.

### Study species

Wild *Eristalis tenax* (Syrphidae: Diptera) individuals, along with some flowers, were collected while feeding from flowers or manure pits at each site and transferred into an insect cage to acclimatize until the experiment. Flies were later released back in the same site immediately after the experiment except those used for cardiac physiology experiments. Only healthy individuals of both sexes, characterized as flying or actively walking with no apparent injuries, were used for the experiments.

### Measurement of antennal sensitivity and threshold for stimulus detection

We standardized a portable electrophysiology (pEAG) rig to perform electroantennograms to measure the antennal response towards the given olfactory stimuli at each site. The basic design is described in Olsson and Hansson (2013) and Batra et al (2019). Briefly, the whole-animal preparation was made, which was kept on a moist paper towel to maintain sufficient humidity in field. Silver (Ag/AgCl) electrodes (via glass micropipettes filled with insect Ringer’s solution) were used as both recording and grounding electrodes. A 600VA UPS (Uninterrupted Power Supply) device was used for supplying power to the airflow stimulator system. The set-up was grounded using an iron rod further connected to the ground (Figure 1). The initial recordings were interrupted by rain and strong winds, so the neurophysiological analyses reported here were performed on the same rig set up in a shelter at each site. Antennal responses of *E. tenax* (reared in lab following Nicholas et al., 2018) were measured in controlled laboratory conditions in Bengaluru as described in Rajan and Mishra et al (2024).

**Figure 1:**
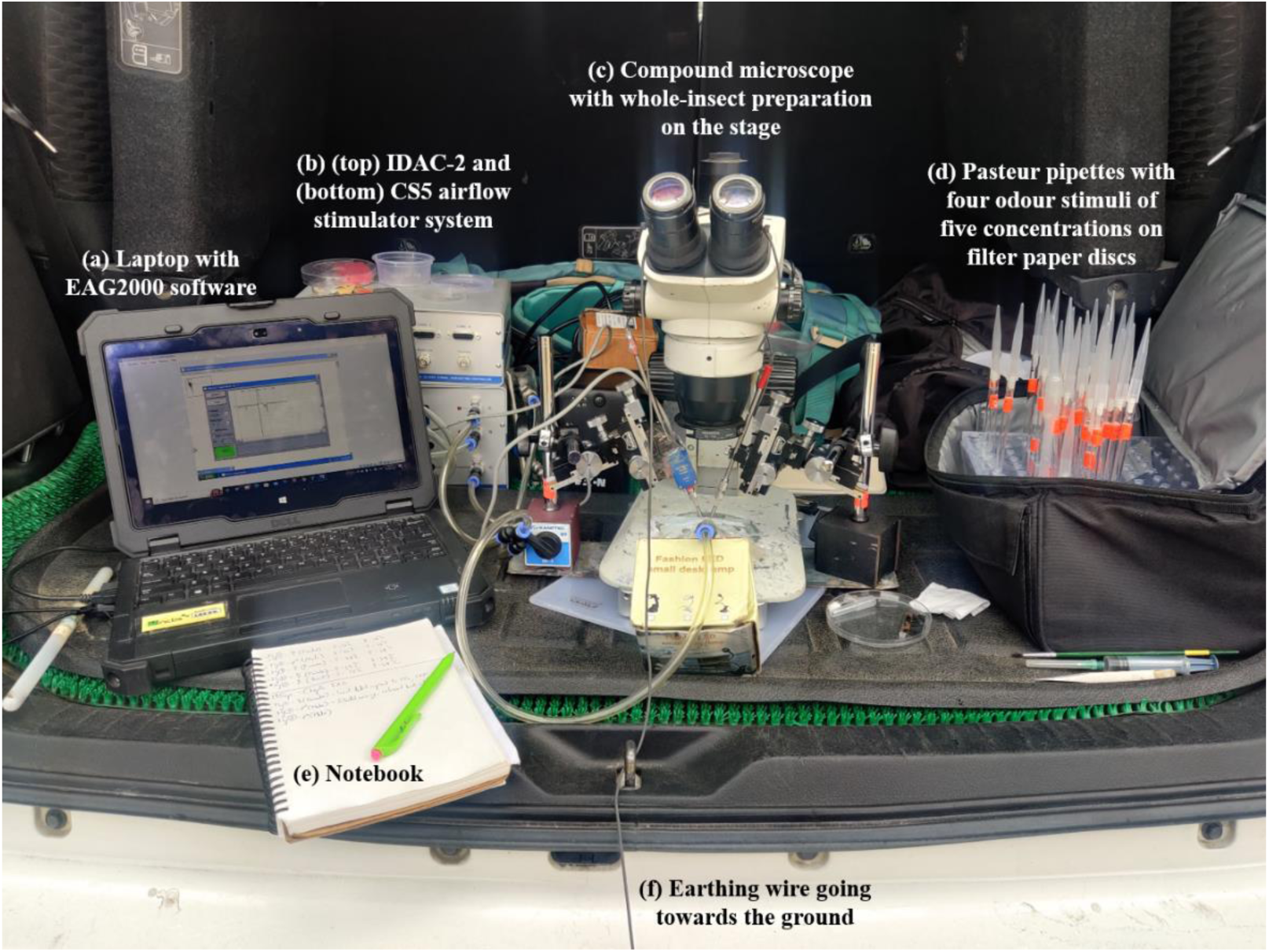
The portable physiology set up to measure the antennal sensitivity and cardiac physiology of *Eristalis tenax* towards common floral volatiles shown in the cargo area of a field vehicle. Micromanipulators and airflow system were removed during cardiac measurements.

Olfactory stimuli were prepared and delivered using previously standardized stimulus delivery set-up (Rajan and Mishra et al., 2024, Tait et al., 2016) with airflow of 2L/min to account for the in-field preparation. The olfactory stimuli were selected from known floral volatiles in the region (Nordström et al., 2017) and prepared using chemicals (>95% purity) purchased from Sigma Aldrich, Bengaluru, India. These consisted of serially diluted doses (10^-1^– 10^-5^) of alpha-pinene, p-cymene, 2-pentylfuran and cis-3-hexenyl acetate dissolved in mineral oil (v/v). The positive control was an odour blend consisting of five common floral chemicals first described by Nordström et al., (2017) and known to elicit antennal response in *E. tenax* (Rajan and Mishra et al., 2024; SI Appendix, S4). A Syntech Intelligent Data Acquisition Controller (IDAC-2) and EAG2000 software were used for data acquisition. Each insect was first given a blank (air), a solvent control (mineral oil) and a positive control stimulus, followed by doses of all chemicals serially from 10^-5^-10^-1^, with a randomized order of chemicals. Only individuals who were tested against all stimuli were used for further analyses. Temperature was measured at the beginning and end of each recording.

### Measurement of heart rate

We performed dissections to record the heart rate at each site using a modified protocol from Thimmegowda et al (2020) in the same setup described for electrophysiology. Briefly, the acclimatized individuals were pinned onto a wax plate and their abdomen dissected under a dissection microscope to expose the beating heart on the dorsal side. A video of the beating heart was recorded for 5 minutes continuously across all 3 abdominal segments, ensuring good resolution for the video analyses (see Video S1: sample heartrate recording (trimmed) of *Eristalis tenax*). Temperature was measured at the beginning and end of each recording.

### Data analyses

During all electroantennographic measurements, the 0.5s odour stimulus was delivered within the first four seconds of starting the recording. Therefore, only the maximum negative deflection values in the first four seconds of the recording were considered. The value for each dose of the volatile was normalized by dividing the maximum deflection value of that volatile with the lowest provided dose (10^-5^). Student t-tests were performed using these normalized values to assess significant differences between the given dose and the lowest dose with the minimum significant dose as the threshold for antennal detection of that volatile. We then performed a 2-way ANOVA to determine the effect of elevation and chemical dose on the antennal responses. We also performed a logistic regression with 95% confidence intervals for each volatile tested with the normalized response at 10^-1^ as the response variable and temperature as the predictor variable. The average heart rate per minute was calculated by manually counting the number of heart beats from the video recordings as described in Thimmegowda et al (2020). Logistic regressions with 95% confidence intervals were then performed with average heart rate per second as the response variable and elevation and temperature as the predictor variables. All data analyses were carried out in RStudio (RStudio 2020) and GraphPad Prism software 8, GraphPad 303 Software. Inc, California, USA.

## RESULTS

We performed electroantennograms on 16 individuals at 3600masl and 19 individuals at 4200masl in Spiti valley, Himachal Pradesh. We performed heart dissection on 67 individuals at 3000masl, 3500masl and 4000masl in two seasons in Lachen valley, Sikkim. We also performed the same experiments with lab-reared individuals of *E. tenax* in controlled laboratory conditions in Bengaluru (900masl), with 52 individuals for EAG and 51 individuals for heart rate measurement.

### Measurement of antennal sensitivity and threshold for stimulus detection

We recorded antennal responses towards five doses of four chemicals at three elevations. We found that the response threshold increased with elevation (Fig. 2, SI Appendix Table S1 (ix)-(xii), (xxi)-(xxiv), (xxxiii)-(xxxvi), (xlv)-(xlviii). We also found that the mean normalised deflection was significantly different across the tested five doses and across three elevations, for all chemicals except for α-pinene (2-pentylfuran: F(2) = 21.821, p = 9.64e-10; α-pinene F(2) = 0.941, p = 0.39114; cis-3-hexenyl acetate: F(2) = 11.708, p = 1.13e-05; p-cymene: F(2) = 5.396, p = 0.00486; 2-way ANOVA followed by Tukey’s HSD test; SI Appendix Table S2). Further, ambient temperature had no effect on the antennal responses of the animals to any volatile (2- pentylfuran: R^2^ = 0.01911, p = 0.2017, alpha-pinene: R^2^ = 0.003005, p = 0.614, cis-3-hexenyl acetate: R^2^ = 0.02103, p = 0.1801, p-cymene: R^2^ = 0.02888, p = 0.1156) (Fig. 2(e)-(h)). Similar results were obtained even when outliers were included in the analyses (SI Appendix, S5). Thus, our results show that the antennal response thresholds to common floral chemicals are reduced with increasing elevation.

**Figure 2:**
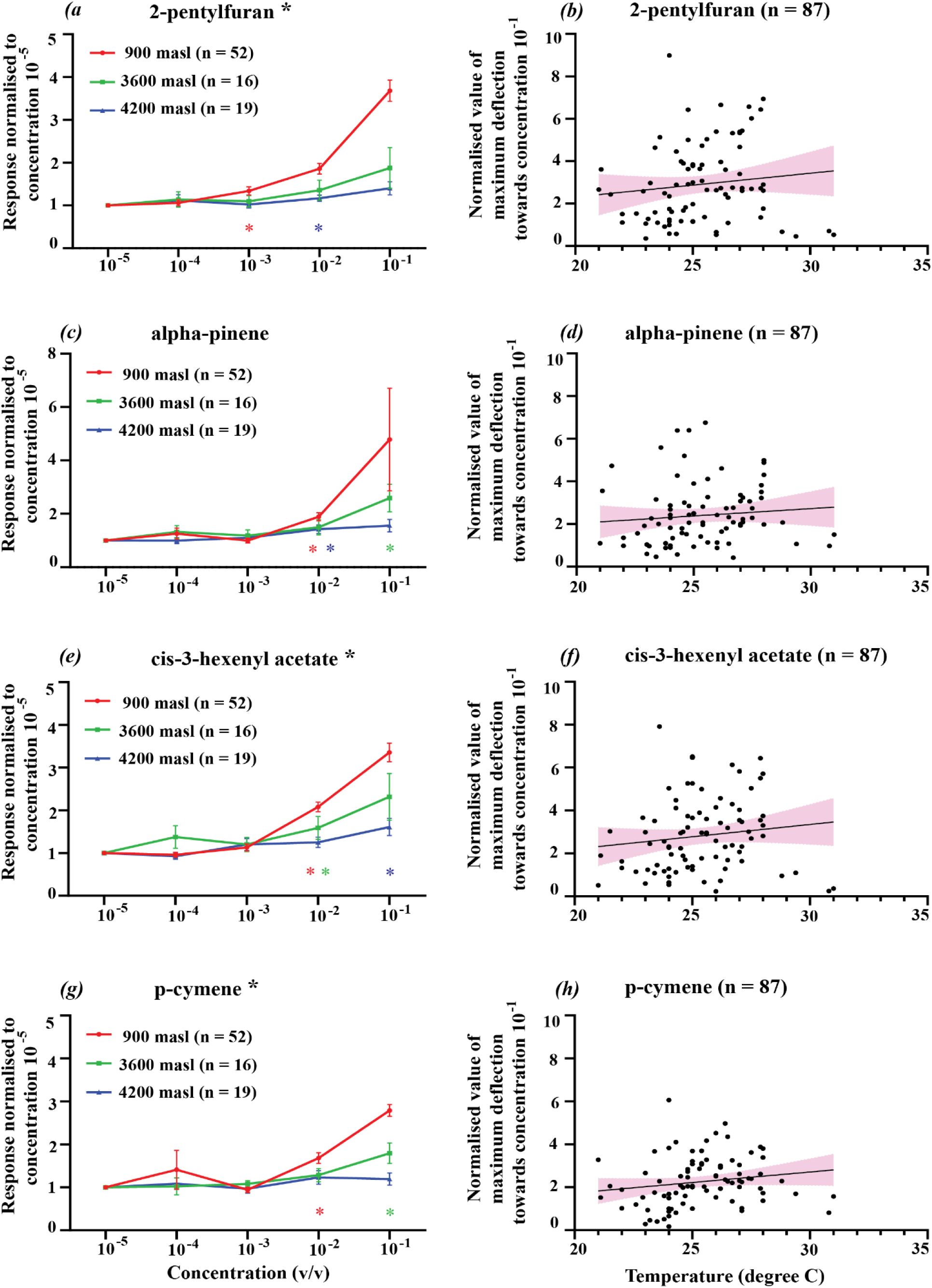
(a)-(d) Antennal responses of *E. tenax* normalised to the lowest dilution (10^-5^) at 900masl (red circles, n = 52), 3600masl (green squares, n = 16) and 4200masl (blue triangles, n = 19) toward four common floral volatiles (a) 2-pentylfuran, (b) alpha-pinene, (c) cis-3-hexenyl acetate, (d) p-cymene. Coloured asterisks indicate the threshold of stimulus detection, based on paired t-test (* for p<0.05), while black asterisks against the chemical names indicate significant differences among the three elevations, based on 2-way ANOVA, Tukey’s HSD post hoc test (* for p<0.05). (e)-(h) Normalised value of maximum deflection towards the 10^-1^ dose plotted against the variation of temperatures for each volatile, where the fitted line is generated by logistic regression, with outliers excluded from the analyses (e) 2-pentylfuran (R^2^ = 0.01465, p = 0.2669), (f) alpha-pinene (R^2^ = 0.009039, p = 0.3867), (g) cis-3-hexenyl acetate (R^2^ = 0.0177, p = 0.2221), (h) p-cymene (R^2^ = 0.02888, p = 0.1156).

### Measurement of heart rate

We found that the average heart rate did not change with elevation (R^2^ = 0.006603, p = 0.5101), but it significantly increased with an increase in local ambient temperature (R^2^ = 0.1437, p = 0.0014) (Figure 3, (a)-(b)). We found the same trend upon comparing these results from field experiments with those obtained from lab-reared individuals in Bengaluru, with a significant correlation between the average heart rate and ambient temperature ((R^2^ = 0.04078, p = 0.02763) (Figure 3, (c)-(d)). Thus, ambient temperature was found to have the most significant impact on average heart rate of insects, irrespective of site, elevation or origin of the individuals.

**Figure 3:**
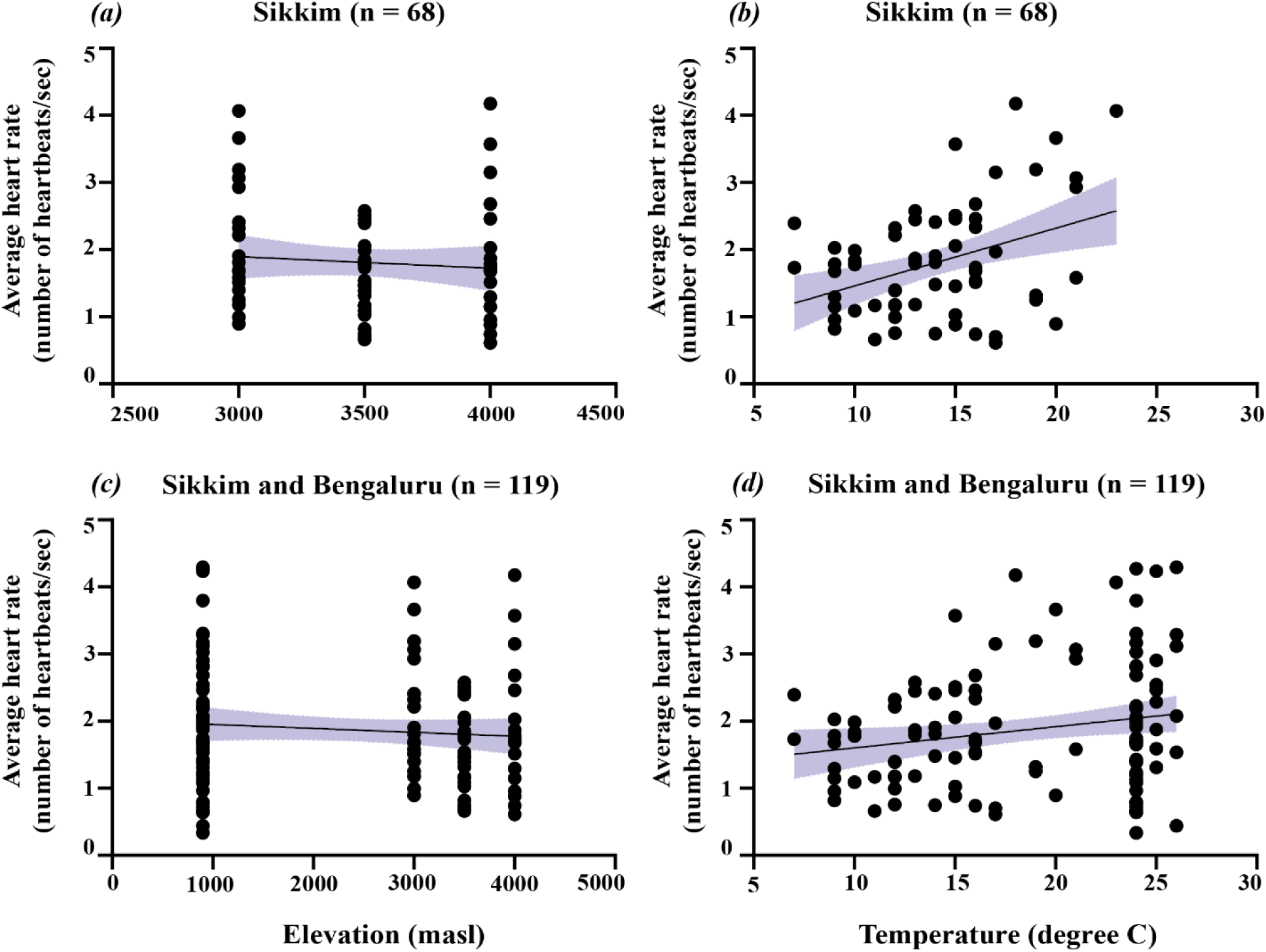
Plots showing variation of average heart rate of *E. tenax* in Sikkim (n = 68) (a) with elevation, (b) with temperature; and when pooled with lab experiments in Bangalore (n = 119), (c) with elevation, and (d) temperature. The fitted line is generated by logistic regression with shaded areas showing 95% confidence intervals.

## DISCUSSION

We report one of the first *in-situ* physiology experiments on wild-caught insects across different elevations in Himalayan mountains. We standardised a portable physiological setup to assess antennal sensitivity and heart rate of *Eristalis tenax* hoverflies using whole insect preparations. Our study showed that the antennal sensitivity of *Eristalis tenax* decreased at higher elevations. Conversely, average heart rate increased with an increase in temperature, but was not affected by elevation. These results show that both elevation and temperature have different impacts on the sensory and cardiac physiology of *E. tenax*.

Our results are consistent with studies showing that average heart rate of *E. tenax* increases with an increase in temperature (Richards, 1963; Lagerspetz and Perttunen, 1962; Andersen et al., 2015). Given that elevation does not have an impact on heart rate, our results show that insects could be able to buffer against hypoxia better than changing ambient temperature. This might have serious implications on their foraging efficiency and ability to obtain sufficient nutrition at higher temperatures. Further experiments involving translocation of insects and transcriptomic profiling can be performed to understand the nuances of their physiological responses.

To our knowledge, this is the first study to perform *in situ* electrophysiology recordings at high elevations. We retained the basic blueprint of a typical laboratory rig (Olsson and Hansson, 2013), with minor additions (i.e. moist tissue under the insect preparation, high airflow speed of 2L/min in a sheltered area) to ensure enough humidity and airflow for the animal. Olfactory sensitivity of humans has been shown to reduce at higher elevations (Burdack-Freitag et., 2011; Altundağ et al., 2014). Our results show the same trend for insects, where the antennal (olfactory) sensitivity of *E. tenax* towards common floral chemicals decreased with elevation. Lack of sensitivity to lower concentrations of chemicals at higher elevations could lead to failure in detecting relevant floral olfactory cues and sensory mismatches, especially after upslope range shifts.

Our results show that abiotic factors have contrasting effects on physiological parameters. While elevation did not impact hoverfly heart rate, changes in elevation significantly altered antennal sensitivity. In contrast, increased temperatures had a significant impact on heart rate, but not antennal response. Our results suggest that as insects shift their ranges to higher elevations with lower temperatures, cardiac physiology might be maintained, but insects might be less efficient at detecting olfactory cues, potentially leading to increased foraging times and reduced foraging efficiencies. At the same time, insects at lower altitudes will experience higher heart rates as temperatures rise, which could impact their survival. Finally, our results provide necessary baseline data (Spence and Tingley, 2020; Shah et al., 2020) about the current physiological status of hoverflies in the tropical Himalayan mountains. These data will be critical not only to understand the survival of insect pollinators in the dynamic high-altitude, montane ecosystem, but also help to predict impacts of climate change events on the survival of future populations.

## ACKNOWLEDGMENTS

The authors thank Dr. Kulbhushansingh Suryawanshi, Deepshikha Sharma, Kalzang Gurmet from Nature Conservation Foundation, Mysuru for assistance in field work. The authors would like to acknowledge Aditi Mishra, Hinal Kharva and Mayuresh Gangal for insightful discussions on the manuscript. We thank the people of Sikkim and Spiti valley (especially teams from field stations of the High Altitude Program of Nature Conservation Foundation) for logistics support and hospitality during the field experiments as well as the Sikkim Forest Department, Himachal Pradesh Forest Department, Pipon and Lachen Dzumsa, Sikkim Police Department, Indo-Tibetan Border Police, Indian Army, Home Department for permits and support.

## ARTIFICIAL INTELLIGENCE (AI) DECLARATION

The authors have not used any Artificial intelligence (AI) tools to generate any part of the manuscript.

## ETHICAL CLEARANCE STATEMENT

We obtained permits from the Department of Forest and Environment, Government of Sikkim (2021-2022: F.no. 78/GOS/FEWMD/BDR/PCCF/Secy/R&E/190 and F.no. 78/GOS/FEWMD/BDR/PCCF/Secy/R&E/146) and the Himachal Pradesh Forest Department (2023: No.WLM-08/Research Study/7729). We also received permission from respective land-owners to perform experiments at the field sites.

