## Supplementary Information for "Surviving in the mountains: Temperature and elevation have contrasting physiological effects on the hoverfly *Eristalis tenax* in the Himalayas"

**TITLE:**

**ABSTRACT:**

Insect populations are experiencing a global decline due to a variety of human-linked environmental changes. Among these changes, how insects' physiology might be affected by predicted upslope migration due to climate change is unknown. Being ectotherms, insect physiology is impacted by abiotic factors like ambient temperature that change with elevation. Here, we performed *in situ* experiments to assess the sensory and cardiac physiology of an important generalist pollinating hoverfly *Eristalis tenax* (Diptera: Syrphidae), across different elevations in the eco-sensitive and biodiverse Himalayan mountains. We built a portable physiology setup and measured hoverfly antennal responses towards common floral volatiles at 3600 masl and 4200 masl. We also recorded their heart rate at 3000 masl, 3500 masl and 4000 masl. We report the first *in situ* physiology experiments performed in the high-altitude Himalayas. Our results show a contrasting impact of elevation and temperature on the sensory and cardiac physiology of hoverflies, with antennal sensitivity decreasing with increasing elevation, while average heart rate increased with temperature, independent of elevation. With upslope migration and climate warming, consequent sensory mismatches and cardiac stress could have deleterious effects on the health of both hoverflies and the vulnerable Himalayan ecosystem.

**S1.** Student t-tests assessing threshold antennal sensitivities of four floral chemicals across three elevations. Mean maximum deflections of a given dose of a tested volatile were compared to the lowest dose ($10^{-5}$). A statistically significant value indicates that the antennal response obtained for that dose was significantly different from the lowest dose, and that dose is classified as the threshold of antennal detection for that chemical at that elevation.

| S.no. | Site | Elevation (masl) | Floral chemical/ olfactory cue | Comparison (concentration pair) | No. of pairs compared | t-value | p-value | Threshold of significant detection |
|---|---|---|---|---|---|---|---|---|
| (i) | Bengaluru | 900 | 2-pentylfuran | $10^{-5}$ and $10^{-4}$ | 52 | 0.8335 | 0.4085 | $10^{-3}$ |
| (ii) | Bengaluru | 900 | 2-pentylfuran | $10^{-5}$ and $10^{-3}$ | 52 | 3.536 | **<0.0001** | |
| (iii) | Bengaluru | 900 | 2-pentylfuran | $10^{-5}$ and $10^{-2}$ | 52 | 6.765 | **<0.0001** | |
| (iv) | Bengaluru | 900 | 2-pentylfuran | $10^{-5}$ and $10^{-1}$ | 52 | 10.72 | **<0.0001** | |
| (v) | Kaza | 3600 | 2-pentylfuran | $10^{-5}$ and $10^{-4}$ | 16 | 0.7645 | 0.4564 | none |
| (vi) | Kaza | 3600 | 2-pentylfuran | $10^{-5}$ and $10^{-3}$ | 16 | 0.7222 | 0.4813 | |
| (vii) | Kaza | 3600 | 2-pentylfuran | $10^{-5}$ and $10^{-2}$ | 16 | 1.527 | 0.1476 | |
| (viii) | Kaza | 3600 | 2-pentylfuran | $10^{-5}$ and $10^{-1}$ | 16 | 1.837 | 0.0862 | |
| (ix) | Kibber | 4200 | 2-pentylfuran | $10^{-5}$ and $10^{-4}$ | 19 | 0.8696 | 0.3959 | $10^{-2}$ |
| (x) | Kibber | 4200 | 2-pentylfuran | $10^{-5}$ and $10^{-3}$ | 19 | 0.2476 | 0.8072 | |
| (xi) | Kibber | 4200 | 2-pentylfuran | $10^{-5}$ and $10^{-2}$ | 19 | 2.39 | **0.028** | |
| (xii) | Kibber | 4200 | 2-pentylfuran | $10^{-5}$ and $10^{-1}$ | 19 | 2.573 | **0.0192** | |
| (xiii) | Bengaluru | 900 | alpha-pinene | $10^{-5}$ and $10^{-4}$ | 52 | 1.243 | 0.2197 | $10^{-2}$ |
| (xiv) | Bengaluru | 900 | alpha-pinene | $10^{-5}$ and $10^{-3}$ | 52 | 0.06721 | 0.946 | |
| (xv) | Bengaluru | 900 | alpha-pinene | $10^{-5}$ and $10^{-2}$ | 52 | 5.583 | **<0.0001** | |
| (xvi) | Bengaluru | 900 | alpha-pinene | $10^{-5}$ and $10^{-1}$ | 52 | 1.969 | **0.0544** | |
| (xvii) | Kaza | 3600 | alpha-pinene | $10^{-5}$ and $10^{-4}$ | 16 | 1.359 | 0.1943 | $10^{-1}$ |
| (xviii) | Kaza | 3600 | alpha-pinene | $10^{-5}$ and $10^{-3}$ | 16 | 0.8443 | 0.4118 | |
| (xix) | Kaza | 3600 | alpha-pinene | $10^{-5}$ and $10^{-2}$ | 16 | 1.742 | 0.102 | |
| (xx) | Kaza | 3600 | alpha-pinene | $10^{-5}$ and $10^{-1}$ | 16 | 3.082 | **0.0076** | |
| (xxi) | Kibber | 4200 | alpha-pinene | $10^{-5}$ and $10^{-4}$ | 19 | 0.0489 | 0.9615 | $10^{-2}$ |
| (xxi) | Kibber | 4200 | alpha-pinene | $10^{-5}$ and $10^{-3}$ | 19 | 0.7651 | 0.4541 | |
| (xxiii) | Kibber | 4200 | alpha-pinene | $10^{-5}$ and $10^{-2}$ | 19 | 2.626 | **0.0171** | |
| (xxiv) | Kibber | 4200 | alpha-pinene | $10^{-5}$ and $10^{-1}$ | 19 | 2.394 | **0.0278** | |
| (xxv) | Bengaluru | 900 | cis-3-hexenyl acetate | $10^{-5}$ and $10^{-4}$ | 52 | 0.6673 | 0.5076 | $10^{-2}$ |
| (xxvi) | Bengaluru | 900 | cis-3-hexenyl acetate | $10^{-5}$ and $10^{-3}$ | 52 | 1.549 | 0.1277 | |
| (xxvii) | Bengaluru | 900 | cis-3-hexenyl acetate | $10^{-5}$ and $10^{-2}$ | 52 | 9.642 | **<0.0001** | |
| (xxviii) | Bengaluru | 900 | cis-3-hexenyl acetate | $10^{-5}$ and $10^{-1}$ | 52 | 10.79 | **<0.0001** | |

| | | | | | | | | |
|---|---|---|---|---|---|---|---|---|
| (xxix) | Kaza | 3600 | cis-3-hexenyl acetate | $10^{-5}$ and $10^{-4}$ | 16 | 1.442 | 0.1699 | $10^{-2}$ |
| (xxx) | Kaza | 3600 | cis-3-hexenyl acetate | $10^{-5}$ and $10^{-3}$ | 16 | 1.223 | 0.2402 | |
| (xxxi) | Kaza | 3600 | cis-3-hexenyl acetate | $10^{-5}$ and $10^{-2}$ | 16 | 2.209 | **0.0432** | |
| (xxxii) | Kaza | 3600 | cis-3-hexenyl acetate | $10^{-5}$ and $10^{-1}$ | 16 | 2.406 | **0.0295** | |
| (xxxiii) | Kibber | 4200 | cis-3-hexenyl acetate | $10^{-5}$ and $10^{-4}$ | 19 | 1.024 | 0.3194 | $10^{-1}$ |
| (xxxiv) | Kibber | 4200 | cis-3-hexenyl acetate | $10^{-5}$ and $10^{-3}$ | 19 | 1.48 | 0.156 | |
| (xxxv) | Kibber | 4200 | cis-3-hexenyl acetate | $10^{-5}$ and $10^{-2}$ | 19 | 2.05 | 0.0552 | |
| (xxxvi) | Kibber | 4200 | cis-3-hexenyl acetate | $10^{-5}$ and $10^{-1}$ | 19 | 2.99 | **0.0079** | |
| (xxxvii) | Bengaluru | 900 | p-cymene | $10^{-5}$ and $10^{-4}$ | 52 | 0.9122 | 0.3659 | $10^{-2}$ |
| (xxxviii) | Bengaluru | 900 | p-cymene | $10^{-5}$ and $10^{-3}$ | 52 | 0.8768 | 0.3847 | |
| (xxxix) | Bengaluru | 900 | p-cymene | $10^{-5}$ and $10^{-2}$ | 52 | 5.381 | **<0.0001** | |
| (xl) | Bengaluru | 900 | p-cymene | $10^{-5}$ and $10^{-1}$ | 52 | 12.81 | **<0.0001** | |
| (xli) | Kaza | 3600 | p-cymene | $10^{-5}$ and $10^{-4}$ | 16 | 0.1058 | 0.9171 | $10^{-1}$ |
| (xlii) | Kaza | 3600 | p-cymene | $10^{-5}$ and $10^{-3}$ | 16 | 1.13 | 0.2761 | |
| (xliii) | Kaza | 3600 | p-cymene | $10^{-5}$ and $10^{-2}$ | 16 | 1.866 | 0.0817 | |
| (xliv) | Kaza | 3600 | p-cymene | $10^{-5}$ and $10^{-1}$ | 16 | 3.289 | **0.005** | |
| (xlv) | Kibber | 4200 | p-cymene | $10^{-5}$ and $10^{-4}$ | 19 | 0.6305 | 0.5363 | none |
| (xlvi) | Kibber | 4200 | p-cymene | $10^{-5}$ and $10^{-3}$ | 19 | 0.3144 | 0.7568 | |
| (xlvii) | Kibber | 4200 | p-cymene | $10^{-5}$ and $10^{-2}$ | 19 | 1.433 | 0.169 | |
| (xlviii) | Kibber | 4200 | p-cymene | $10^{-5}$ and $10^{-1}$ | 19 | 1.373 | 0.1868 | |

**S2.** 2-way Analysis of Variances (ANOVAs) comparing antennal sensitivities across three elevations of four chemicals in five doses. Elevation and dose of chemical were predictor variables and the normalized value of the maximum deflection was the response variable. The F-statistic, the corresponding p-value and degrees of freedom are reported, along with the pairs showing significant differences in the group means obtained from post-hoc Tukey’s Honest Significant Difference (HSD).

| S. no. | Chemical | Comparison | F-value | p-value | Significant pairs (with post-hoc Tukey's HSD) in decreasing order of effect size | Degrees of freedom |
|---|---|---|---|---|---|---|
| 1. | 2-pentylfuran | Across doses | 61.54 | **< 2e-16** | Dose: $10^{-1}$ and $10^{-5}$, $10^{-1}$ and $10^{-4}$, $10^{-1}$ and $10^{-3}$; Elevation: 900-4200 masl, 3600-4200 masl | 4 |
| | | Across elevations | 21.821 | **9.64e-10** | | 2 |
| | | Dose*Elevation | 9.923 | **1.12e-12** | | 8 |
| 2. | α-pinene | Across doses | 4.546 | **0.00133** | Dose: $10^{-1}$ and $10^{-5}$, $10^{-1}$ and $10^{-3}$, $10^{-1}$ and $10^{-4}$ | 4 |
| | | Across elevations | 0.941 | 0.39114 | | 2 |
| | | Dose*Elevation | 0.663 | 0.72381 | | 8 |
| 3. | cis-3-hexenyl acetate | Across doses | 64.49 | **< 2e-16** | Dose: $10^{-1}$ and $10^{-5}$, $10^{-1}$ and $10^{-4}$, $10^{-1}$ and $10^{-3}$; Elevation: 900-4200 masl | 4 |
| | | Across elevations | 11.708 | **1.13e-05** | | 2 |
| | | Dose*Elevation | 6.655 | **3.30e-08** | | 8 |
| 4. | p-cymene | Across doses | 14.406 | **4.97e-11** | Dose: $10^{-1}$ and $10^{-5}$, $10^{-1}$ and $10^{-3}$, $10^{-1}$ and $10^{-4}$; Elevation: 900-4200 masl | 4 |
| | | Across elevations | 5.396 | **0.00486** | | 2 |
| | | Dose*Elevation | 2.121 | **0.03284** | | 8 |

**S3.** Seasonal variation in the average heart rate of *E. tenax* at 3000, 3500 and 4000 masl at Lachen valley, Sikkim with respect to (a) elevation ($R^2 = 0.2526$, $p = 0.3931$) and (b) temperature ($R^2 = 0.05189$, $p = 0.2178$) in May; (c) elevation ($R^2 = 0.007092$, $p = 0.6202$) and (d) temperature ($R^2 = 0.2341$, $p = 0.0024$) in September. The same experiment was repeated at 3600 masl and 4200 masl in Spiti valley, Himachal Pradesh, showing variations in average heart rate of *E. tenax* with respect to (e) elevation ($R^2 = 0.01979$, $p = 0.5777$) and (f) temperature ($R^2 = 0.02580$, $p = 0.5243$). The fitted line is generated by logistic regression with shaded areas showing 95% confidence intervals. The statistical differences between seasons could be because the temperatures in Sikkim have different variation (7°C in May vs. 12°C in September) and are significantly different ($t = 4.8266$, $p < 0.0001$) in May vs. September. We were able to better capture the variation in the abiotic conditions in September, while we could sample limited a temperature range in Spiti valley, Himachal Pradesh.

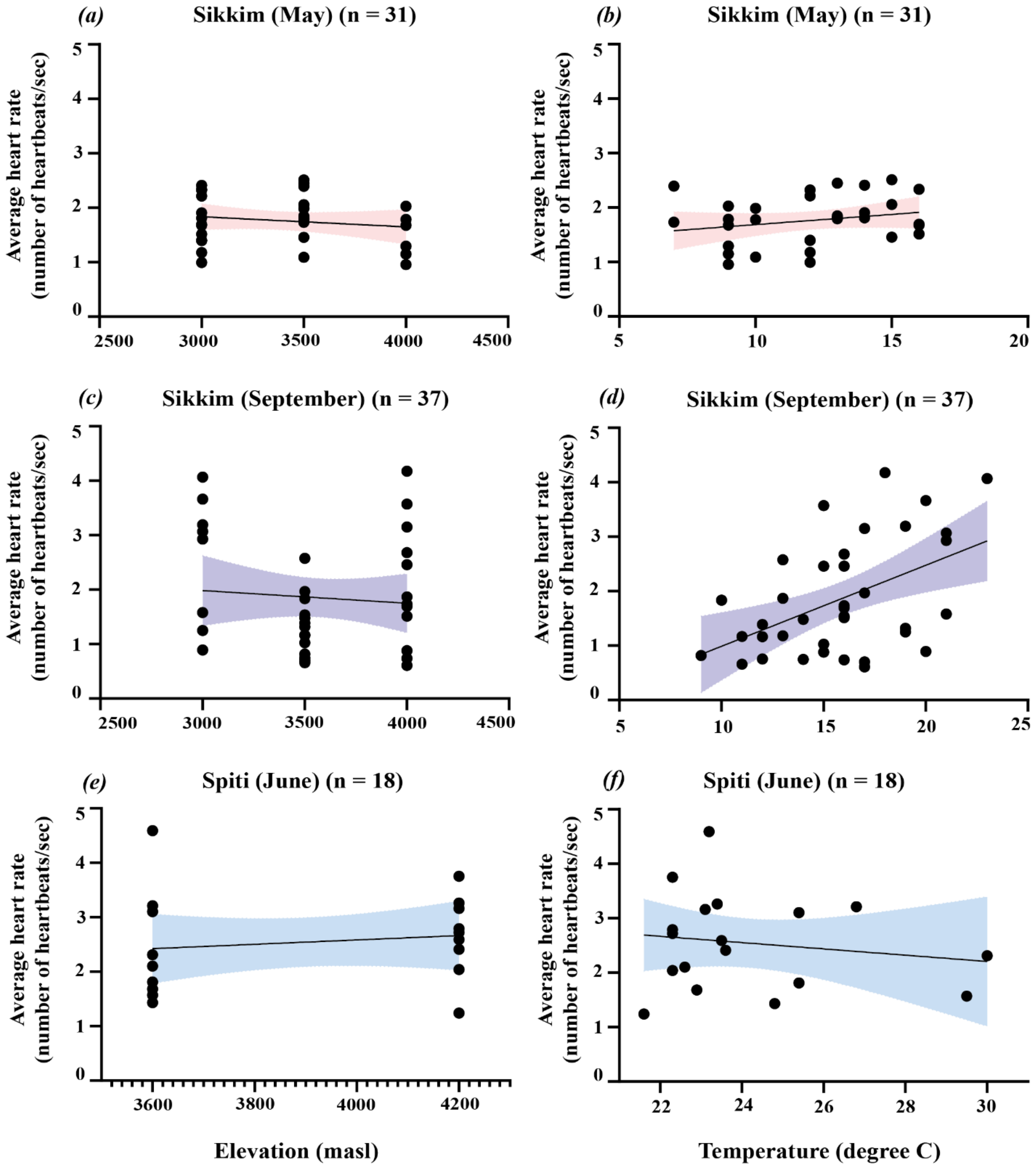


**S4.** Recipe for positive control olfactory blend (from Nordström et al., 2017):

| Chemical | Amount (µl) |
|---|---|
| 2-ethyl toluene | 16.1 |
| p-cymene | 11.8 |
| R-limonene | 24.1 |
| undecanal | 50 |
| 6-methyl-5-hepten-2-one | 17.3 |
| mineral oil | 82.7 |

**S5.** Normalised values of maximum deflection towards the $10^{-1}$ dose plotted against the variation of temperatures for each volatile, where the fitted line is generated by logistic regression with outliers included in the analyses (a) 2-pentylfuran ($R^2 = 0.01911$, $p = 0.2017$), (b) alpha-pinene ($R^2 = 0.003005$, $p = 0.614$), (c) cis-3-hexenyl acetate ($R^2 = 0.02103$, $p = 0.1801$). Ambient temperature had no significant effect on the antennal responses of the animals to any volatile.

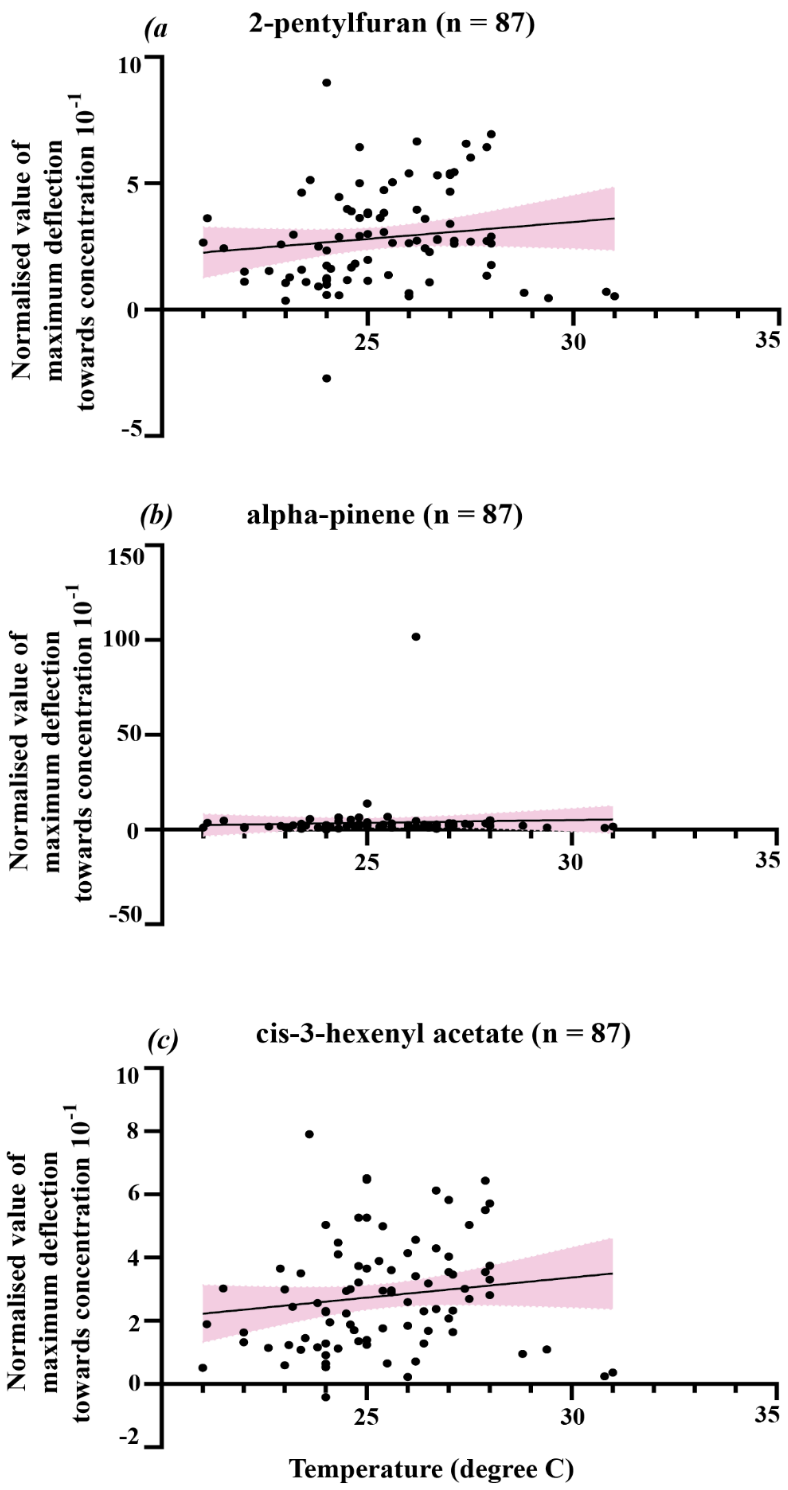


**S6.** Video S1: sample heartrate recording (trimmed) of *Eristalis tenax*
